## Supplementary material for "Why the brown ghost chirps at night"

**Appendix 1: Detecting beats at a distance**

In weakly electric fish, the electric fields generated by electric organs are typically in the range of a few mV, depending on the species. Brown ghosts generate EODs with an amplitude of roughly 5 mV (Rasnow et al., 1993) as measured at the head and tail extremities of the fish body. Considering that the decay of dipole moments is regulated by the following law:

$V=\frac{q}{\varepsilon_{0}}\cdot\frac{s\cdot cos \theta}{4\pi d^{2}}$ **equation (1)**

Where *V* is the voltage measured at a distance *d* (m), *q* is the dipole charge (in Coulombs), *epsilon* is the vacuum dielectric constant, *s* is the distance between the dipole charges (m), and *theta* is the angle of the vector joining the point at distance d and the dipole midpoint (Knudsen, 1975). While this equation is useful to describe the static dependency of field intensity on distance for any measuring point in space, electric fields can also be represented as the vector sum of electric gradients. The formula describing field gradients can be obtained from the formula above by resolving it for each charge in the system with reference to a given point in space moving through it. The field is instantaneously generated by the 2 dipole charges, and so can be described by the formula (Knudsen, 1975):

$E \approx\frac{q}{d^{2}}\cdot\left[ \left( 1-\frac{2s}{d} \right)- \left( 1-\frac{2s}{d} \right) \right]= -\frac{4\cdot q\cdot s}{d^{3}}$ **equation (2)**

Equation (2) clearly shows that within an electric field, the voltage drops with the cube of distance. At double the distance, the field amplitude drop is 8-fold (see Benda, 2020 for a recent and exhaustive description of the physical properties of weakly electric fields). According to these considerations, the range at which brown ghosts can realistically detect other electric fields depends on whether or not active electrolocation is used. In fact, while the dipole moment generated by an EOD alone dissipates according to equation (2), the amplitude modulation (AM) resulting from the interaction of 2 or more EODs depends on the decay constants of all composing signals. Now, it is known that brown ghosts are not capable of detecting single EODs and reading their absolute voltage values, rather they sense the AM generated by the interacting EODs (i.e. the *beat*). From the amplitude of these AMs (i.e. the *beat contrast*), brown ghosts can infer the relative distance, the size and even the rate of motion of an approaching conspecific (Assad, 1997; Babineau et al., 2006; Kelly et al., 2008). The beat contrast can be easily calculated as the ratio of the amplitude of the detected signal and the EOD amplitude of the receiver fish (A_1_/A_0_ x 100). Behavioral experiments have shown that responses to beats can be elicited in different knifefish species at electric field gradients as low as a few μV/cm (0.6 μV/cm in *Eigenmannia virescens*, Kaunzinger and Kramer, 1995; 0.2 μV/cm *Sternopygus macrurus*, Fleishman et al., 1992 and *Apteronotus albifrons*, Knudsen 1974). In *Eigenmannia virescens*, behavioral responses to EOD AMs were observed at 0.02-0.03% beat contrast (Carr et al., 1986; Kawasaki et al., 1997), although other authors reported for the same species slightly higher values (0.16% contrast, Rose and Heiligenberg, 1985). Although beat detection thresholds in *Eigenmannia virescens* and brown ghosts may slightly differ, estimates of field gradient sensitivity have been estimated to be similarly low in brown ghosts (0.1 μV/cm, Rasnow, 1996). Therefore, we could expect similar beat sensitivity levels also in this species. Based on these considerations, we determined the values of EOD amplitude and beat contrast for a reference fish generating an electric field resulting in an average 5 mV voltage measure around its body and a 1.25 mV/cm field gradient starting at 2 cm distance. The average voltage was estimated from skin measurements conducted in brown ghosts and referenced to a ground electrode placed at “infinite” distance” (Rasnow, 1993). At 2 cm distance and starting from 5 mV, the average voltage drops at 1.25 mV (according to equation 1) so choosing this value as a starting point for field measurements at the same distance represents a slight overestimation (the electric field decays with the cube of distance according to in equation 2). Beat contrast is estimated with reference to the first value and only estimates from further distances are shown.

*In the diagram, electric field ranges are represented by the oval shaped lines drawn around the fish and are based on the voltage values obtained using the equations above. Ranges for passive electrolocation (dotted line) are indicated for reference (Lissmann and Machin, 1958; Von der Emde, 1999; Knudsen, 1974). The beat contrast values obtained are plotted as a function of distance. The example illustrates the following situation: a 1.25 mV/cm field is used as reference (on the right) and the beat induced by a neighboring 0.7 mV/cm field at different distances is calculated. The composite signals obtained at representative distances are displayed by the graphs on the top together (blue) with their AM (red). The value in (3) is the lowest possible contrast level at which the moving fish can be detected and corresponds to a limit distance of about 60 cm between the two fish. This estimate is close to the reported range for chirping (32 cm, Henninger et al., 2018; Zupanc et al., 2006; Hupé and Lewis, 2008) and EOD envelope processing (Fotowat et al., 2013).*
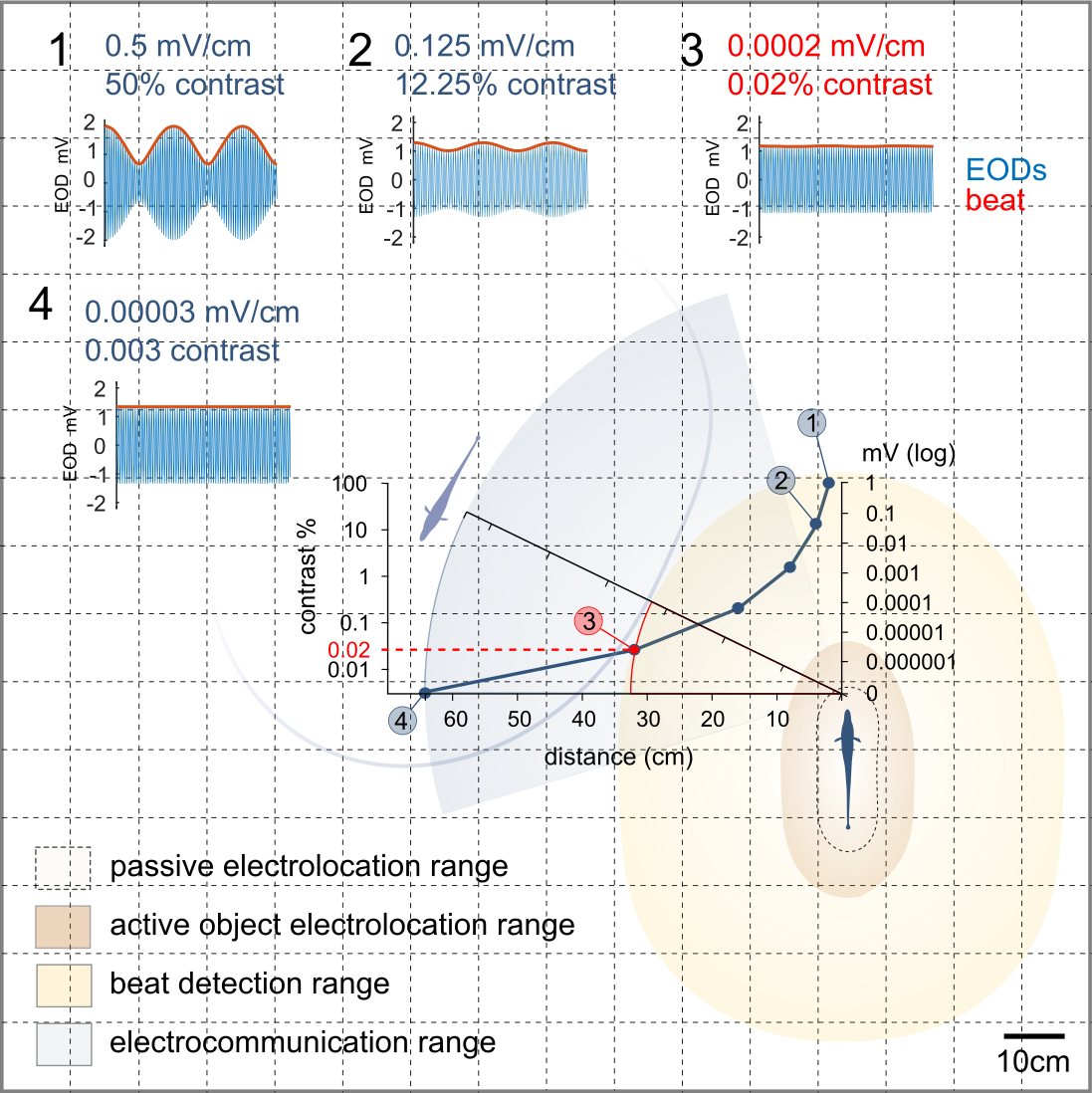

As mentioned, active electrolocation of conspecifics implies the summation of 2 or more EODs and the detection of the resulting beat. Because the maximum amplitude of the resulting composite signals is higher than each component - and because beat contrast is calculated using a different reference depending on the receiver - the threshold for beat detection depends on the decay of the contributing signals and is represented by the minimum AM detectable by each fish (other_EOD/own_EOD*100). For example, if a “fish 1” producing a 1.25 mV EOD swims near a “fish 2” conspecific having a 0.7 mV EOD, when they are 16 cm apart, the maximum AM produced by the 2 EODs can be roughly EOD1+EOD2 = (1.25 + 0.0014) mV, which corresponds to a beat contrast of 0.14 %. For fish 2: EOD2+EOD1 = (0.7 + 0.002) mV, corresponding to a beat contrast of 0,28%. Therefore, at any given distance, different fish will detect beats of different amplitudes. It follows that, if beat detection thresholds can be considered comparable in all brown ghosts, fish with EODs of lower amplitude (usually fish of smaller size such as juveniles or females), will have a wider beat detection range.

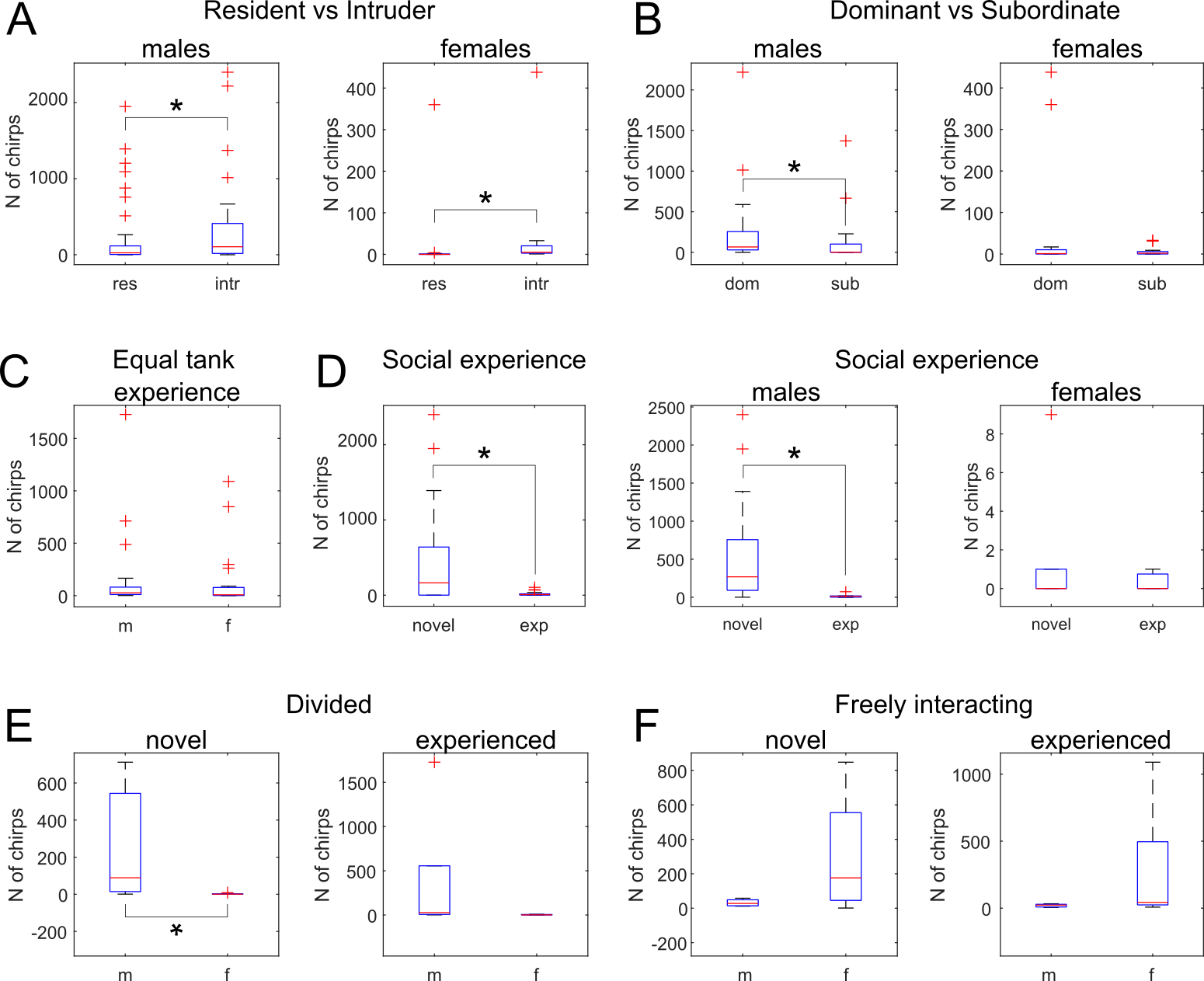
**Figure S1. Chirping in different social contexts and different tank experience conditions.** A: Total chirp counts obtained from resident-intruder assays. Resident fish were housed for 1 week in the test-aquarium while intruders were introduced only at the moment of testing (males, Mann-Whitney U = 0.032, females U < 0.001). B: Total chirp counts recorded during dominant-subordinate interactions (males, Mann-Whitney U = 0.015). C: Total chirp counts obtained from dyadic interactions in which both fish were novel to the test environment. D: Chirp counts evaluated during first time pairing (novel) and after 1 week of pairing (exp, experienced; pooled data, Mann-Whitney U = 0.016; males, Mann-Whitney U = 0.002). E: Chirp counts obtained from opposite sex pairs at the beginning (novel) and at the end of a 4 week-long water conductivity decrease protocol (used to simulate the reproductive season; novel, Mann-Whitney U = 0.034). Fish pairs were interacting only electrically, across a plastic mesh barrier (divided). F: Chirp counts relative to female-male interactions in absence of any mesh barrier (freely interacting). Chirps produced by opposite-sex pairs at the end of the water conductivity changes (experienced) are compared with chirp rates of female-male pairs recorded in absence of prior experience (novel). All chirp counts refer to 1 hour-long recordings except those of dominant-subordinate pairings, which lasted 30 minutes; see methods for details on the behavioral experiments.

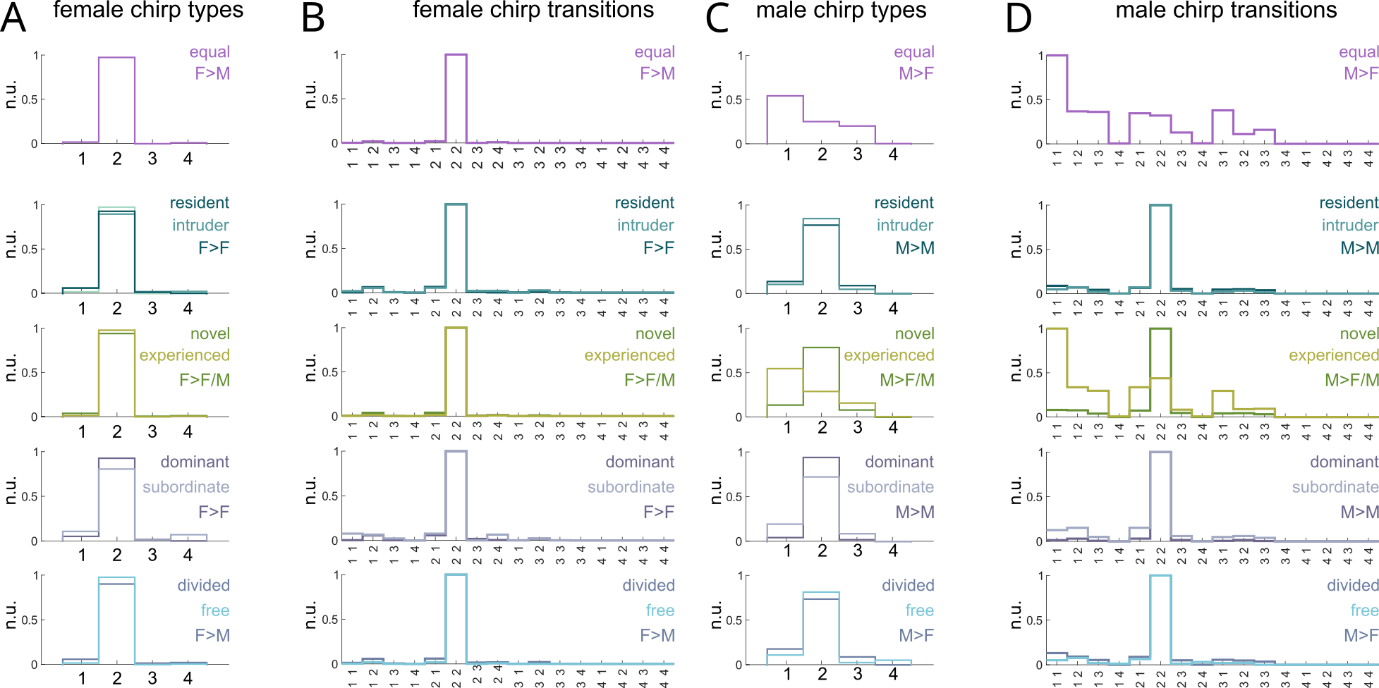

**Figure S2. Effect of social context and environmental experience on chirping behavior**. A: Histograms showing the normalized chirp type distribution relative to female senders and receivers of both sexes (F>F or F>M) in different behavioral contexts. Note the almost identical relative abundance (normalized) of different chirp types. B: Chirp type transitions displayed by female chirpers during different kinds of encounters. In all cases the most common sequence is “type 2-type 2” (the numbers on the X-axis represent the 4 different types, see Figure 1). C: Histograms showing the normalized chirp type distribution for male chirpers. The only difference is observed during male-female interactions in conditions of equal tank experience (“equal”). D: Occurrence of chirp transitions in males is similar to those observed in females with the exception of male-female pairs with equal tank experience or longer-term experience with each other. This is likely due to the lower amount of type 2 chirps produced at lower DFs which in turn results in a relatively higher number of larger chirps (chirp type dependency on DF can be better visualized in Figure 1F and Figure 5C,D).

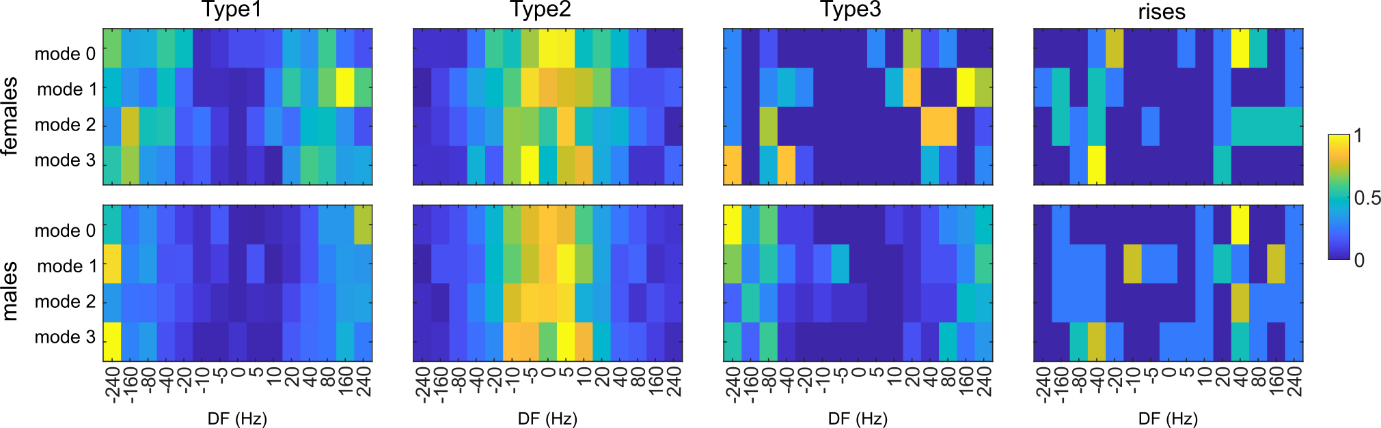

**Figure S3. Sex comparison of fish responses to playback chirps by DF.** A: Heatmaps of responses to different playback signals sorted by DF (X-axis) and type of playback chirps (Y-axis, sine = no chirps, type 1, type 2 and rises). The chirps produced by female and male fish are sorted by type and shown in 4 different maps. Chirp counts are normalized on the max value per type. For a description of playback chirps see methods.

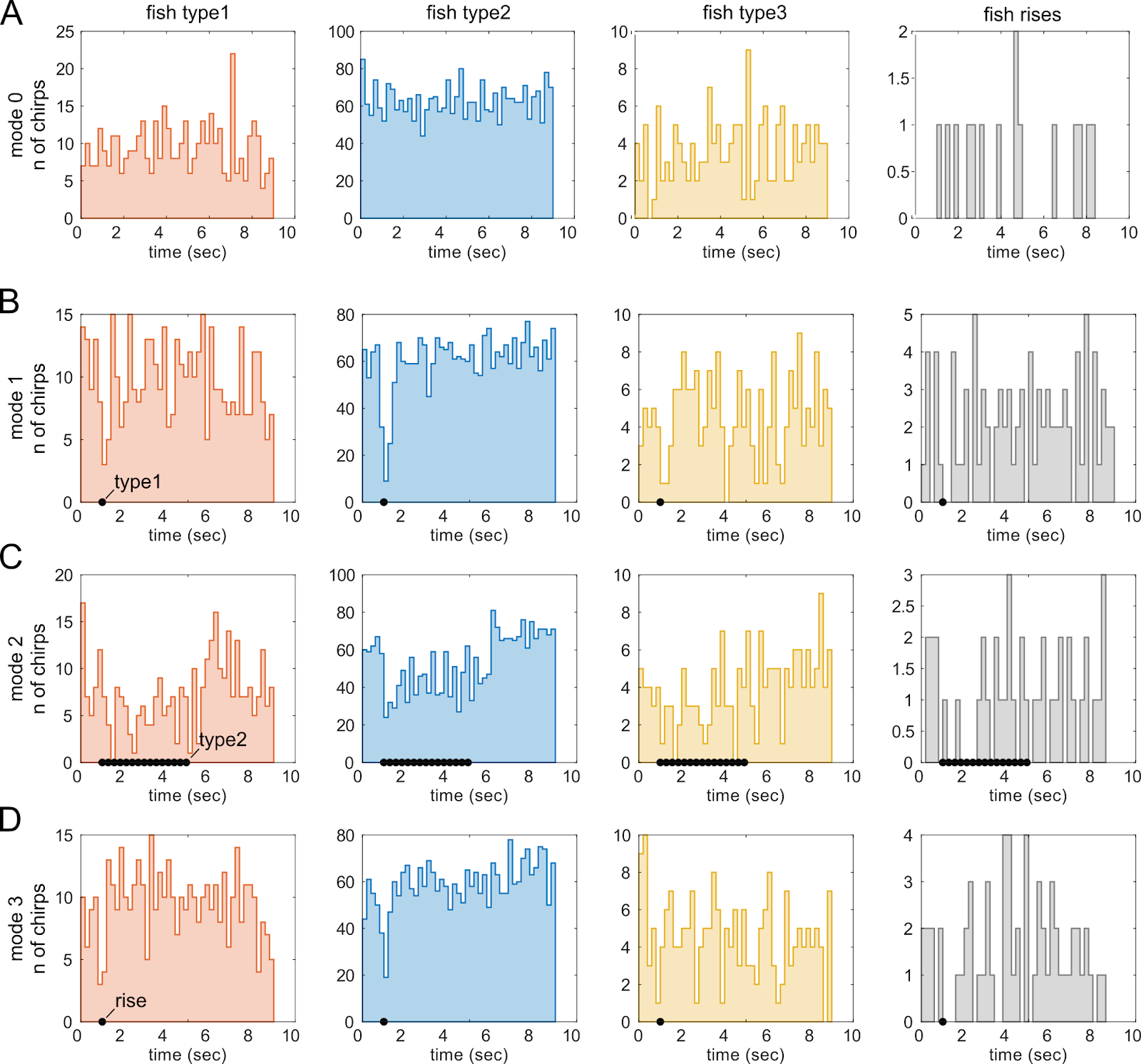

**Figure S4. Playback chirps temporarily suppress chirping in freely swimming fish.** A-D: Peristimulus time-histograms (PSTH) of chirps produced by 16 fish (8 females and 8 males) during playback experiments. Fish responses to different DFs (±240 Hz, ±160 Hz, ±80 Hz, ±40 Hz, ±20 Hz, ±10 Hz, ±5 Hz, 0 Hz) are pooled together. Results for different types of chirps are displayed in different columns while each row is related to chirps produced in response to a given playback mode (mode 0 = plain sinewave, mode 1 = type 1 chirps at 0.2 Hz, mode 2 = type 2 chirps at 3 Hz, mode 3 = rises at 0.2 Hz; see methods for details). The PSTHs show that the main effect of playback chirps (black dots) on the chirps produced by the fish is a brief temporal suppression (accentuated when chirps are repeated in trains, C) but no sign of any significant temporal correlation, except for a transient suppression.

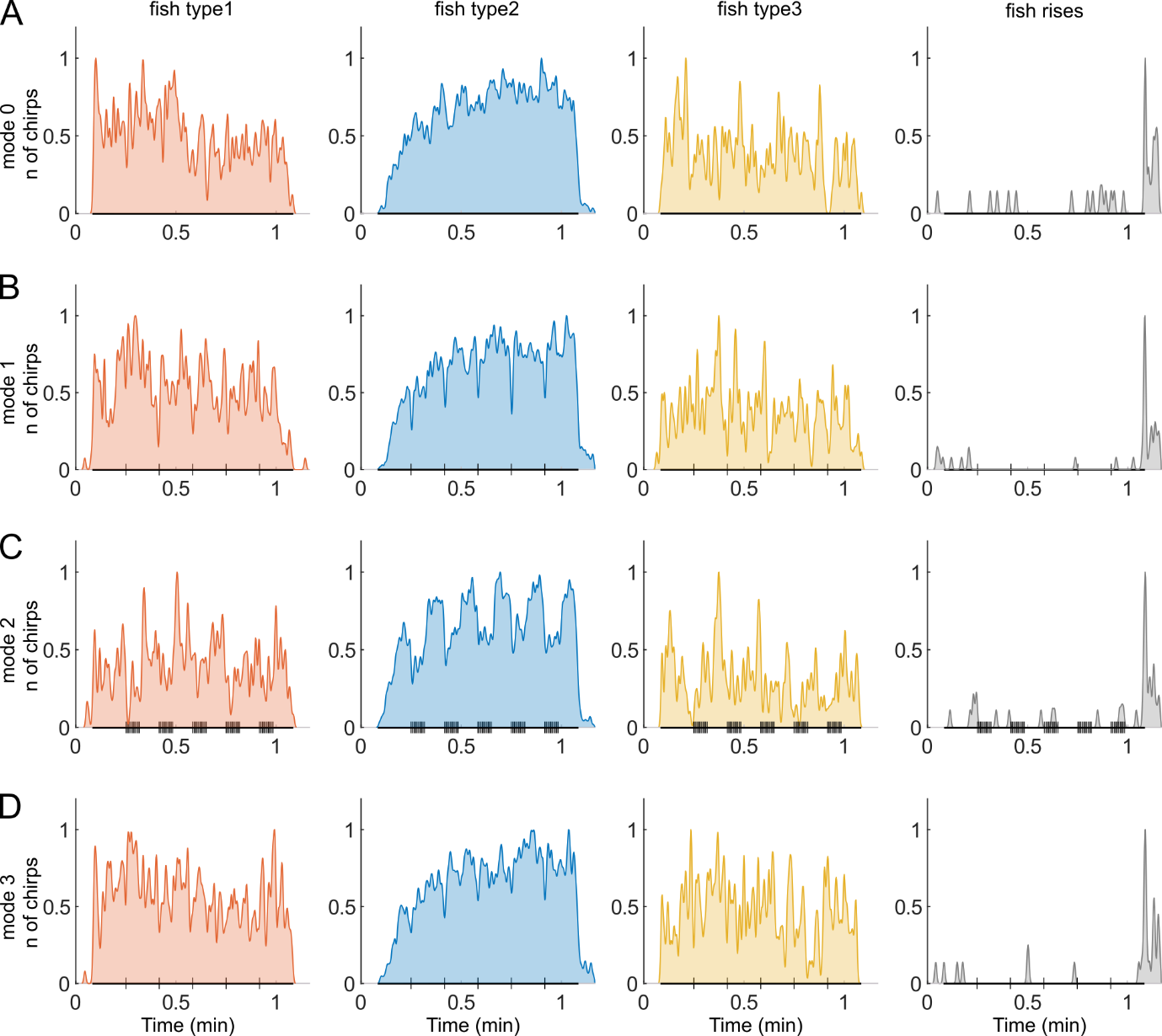

**Figure S5. Chirp responses to playback EODs containing chirps**. A: Average number of chirps of different types (normalized) produced over the course of 1-minute long playback trials (mode 0, sinewave frequency range -240 Hz to +240 Hz). The timing of playback stimuli is represented by the thicker line on the X axis. Vertical ticks on the same axis represent playback chirps. B: Responses to the same set of sine wave EODs to which type 1 chirps were added (mode 1). C: Responses to EODs containing 3Hz trains of type 2 chirps (mode 2). Note the stronger inhibition of fish chirping exerted by trains of type 2 chirps. D: Responses to EODs containing playback rises (mode 3). See method section for details on the playback experiments.

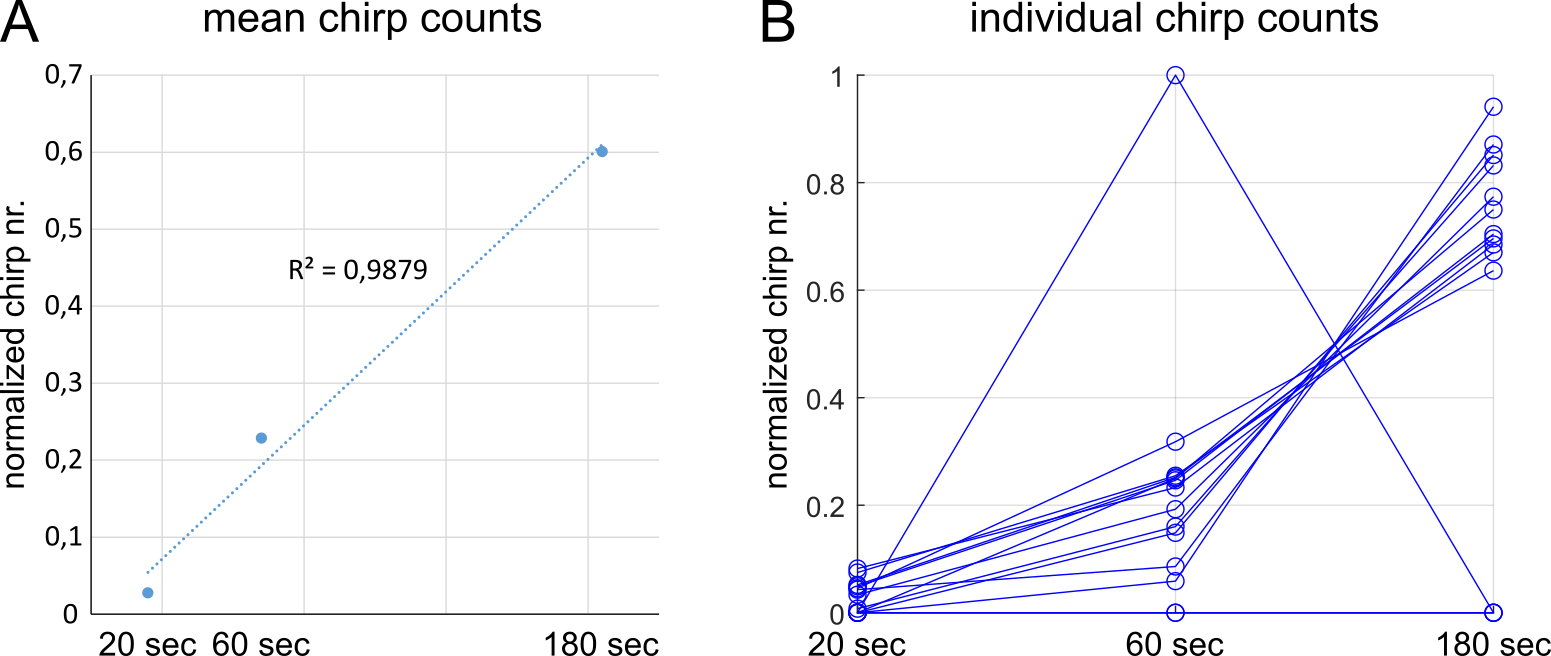

***Figure S6. Linear correlation of chirp counts and playback duration (playback frequency ramps).*** *A: Correlation coefficient of the mean chirp counts (normalized) recorded as a response to playback frequency ramps. B: Line plots related to the individual subjects. The outlier fish emitted 1 chirp during the 60 sec trial and no chirps in the other 2 trials. Fish never producing any chirp were excluded (N=2).*

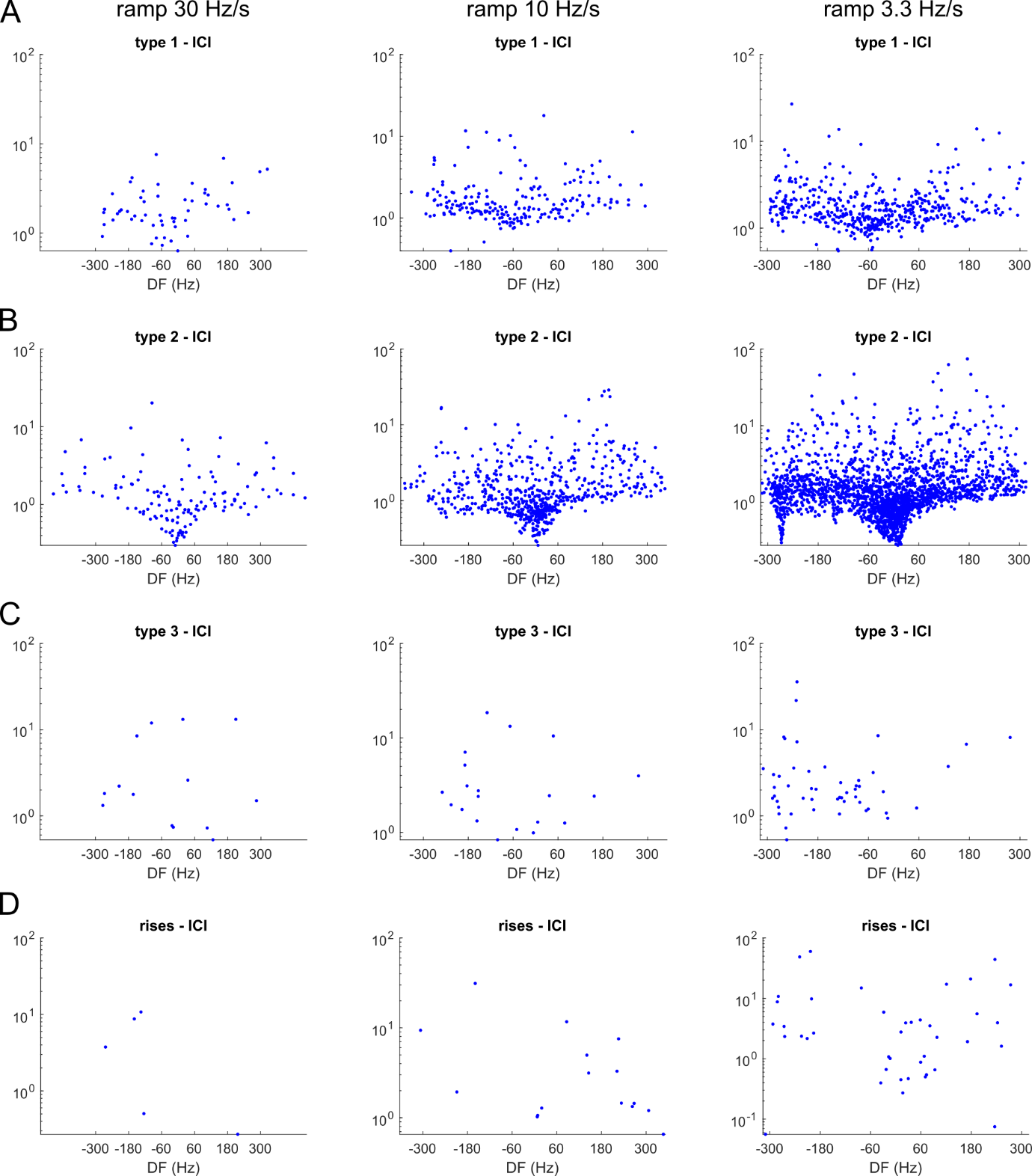

***Figure S7. Frequency ramp playback experiments, inter-chirp intervals (ICI) of different chirp types.*** *A-D) The inter-chirp latencies following each type of chirp are reported for each of the 3 different frequency ramps (see values on the top). These represent the time passing after a given chirp, before a chirp of any given type would follow. Low ICI values are found more often around the DF= 0 Hz (more markedly for type 2 chirps). Low values are also found at trial onset, due to the often higher chirp rates observed at the beginning of a playback trial (see type 1 and type 3 chirps in Figure S5 for instance).*

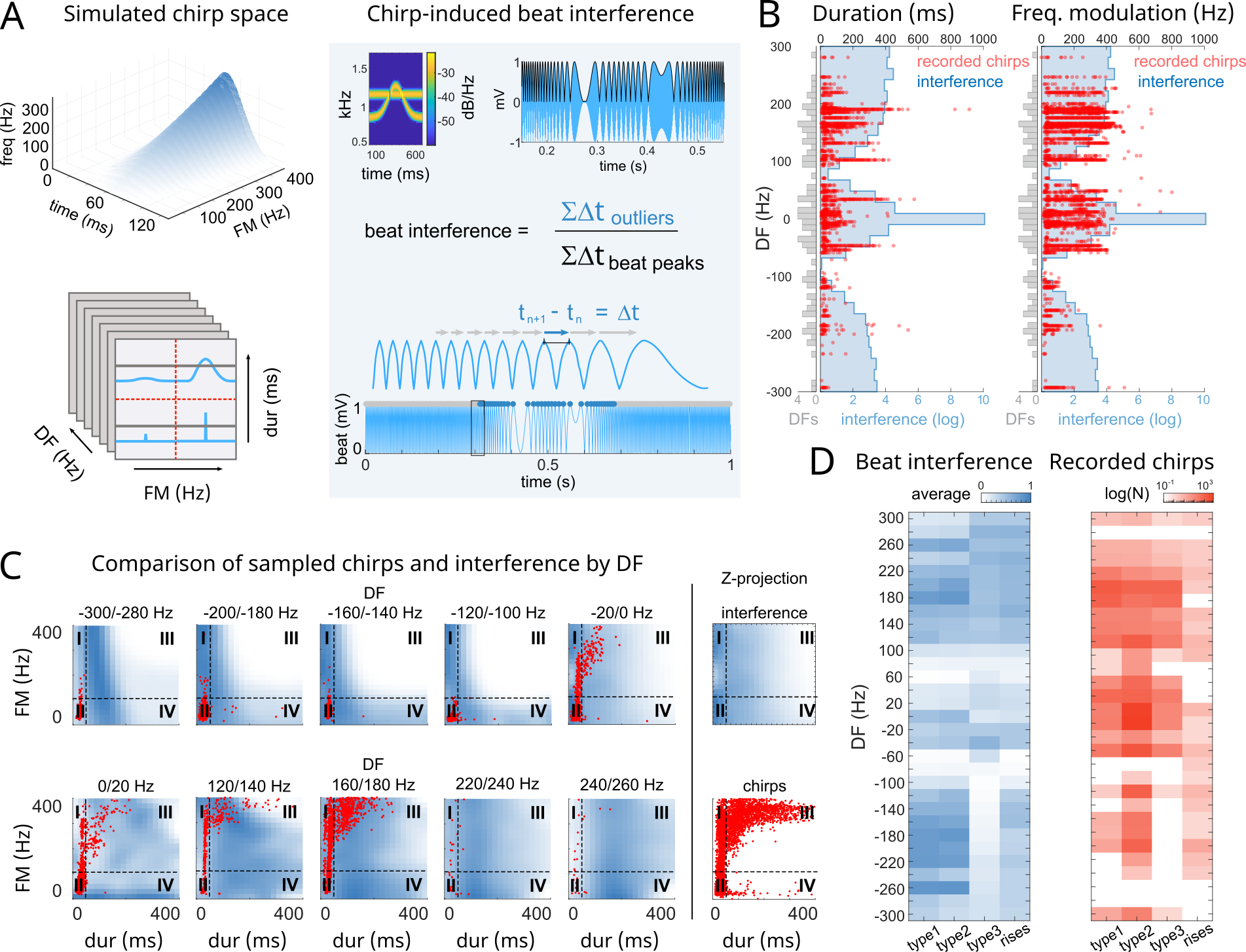

***Figure S8. Chirp interference with beat regularity.*** *A: Artificial chirps were generated on an 862 Hz EOD baseline covering a 0-400 ms duration range and a 0-400 Hz peak amplitude. An additional EOD signal was added to the chirping signal (-300-300 Hz DF range) to simulate a wide range of beat frequencies. The sign of the DF is referred to the baseline reference EOD. The beat interference induced by chirps was calculated as the ratio of the cumulative duration of beat interpeak intervals (IPI) affected by a chirp (i.e. outliers in the IPI population relative to each EOD pair) on the total cumulative beat IPI duration (including outliers) within a 700 ms time window (see methods for details). B: Duration and frequency modulation (FM) histograms of recorded chirps (red, N = 30486) sorted by DF and matched to their estimated beat interference (blue). The gray histograms on the Y-axis represent the beat frequencies sampled. C: Normalized heatmaps showing examples of chirp-induced beat interferences (color coded in blue) calculated at different DF values. Real chirps produced at the same DF are overlaid in red. Only overlapping FM and duration ranges are shown. In the plots on the extreme right: beat interference values are summed over all DFs (top) and are shown next to the corresponding chirps (bottom). D: Comparison of average chirp-induced beat interferences sorted by type (left map, blue) and the actual observed chirps (log N, right map in red).*

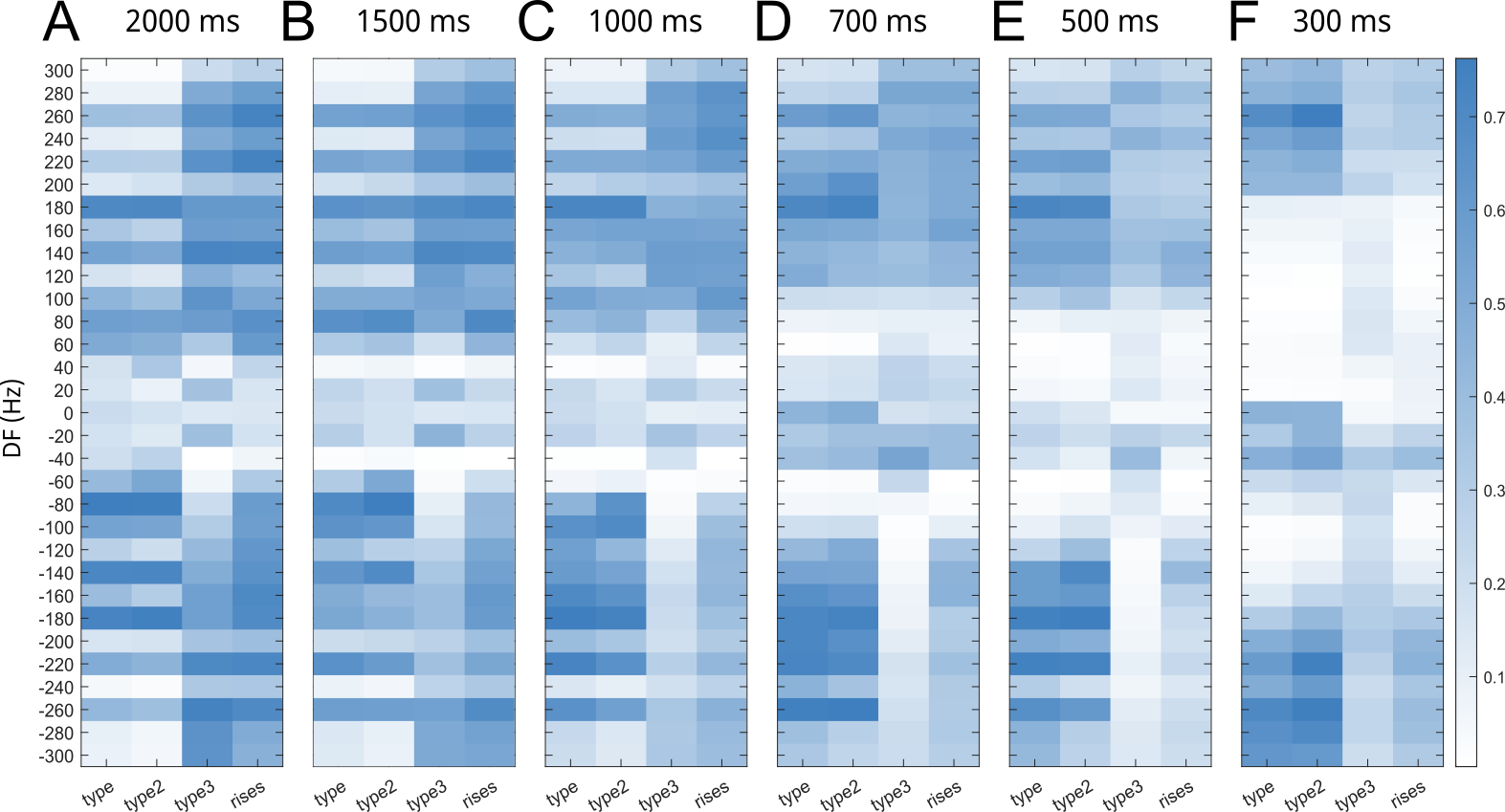

***Figure S9. Beat interference calculated using different time-windows.*** *The effect of the size of the analysis window on the beat interference estimates for the different types of chirps can be evaluated by comparing the heatmaps above. Since the estimate depends on the number of outliers among the beat cycles included within a fixed time interval, the interference value will decrease/change depending on the number of beat cycles considered (i.e. the size of the analysis window): a small window size will have more outliers for fewer beat cycles but a single chirp will affect a larger number of cycles (F). When a larger analysis window is used (A-C), the effect at low DFs is diluted as fewer cycles are affected overall. Other factors potentially affecting the interference estimate include: the algorithm used to detect outliers (here, the outliers are detected in the upper and lower quartiles in the distribution of peak durations) and the phase at which the chirp occurs (here the average of 4 phases is used).*

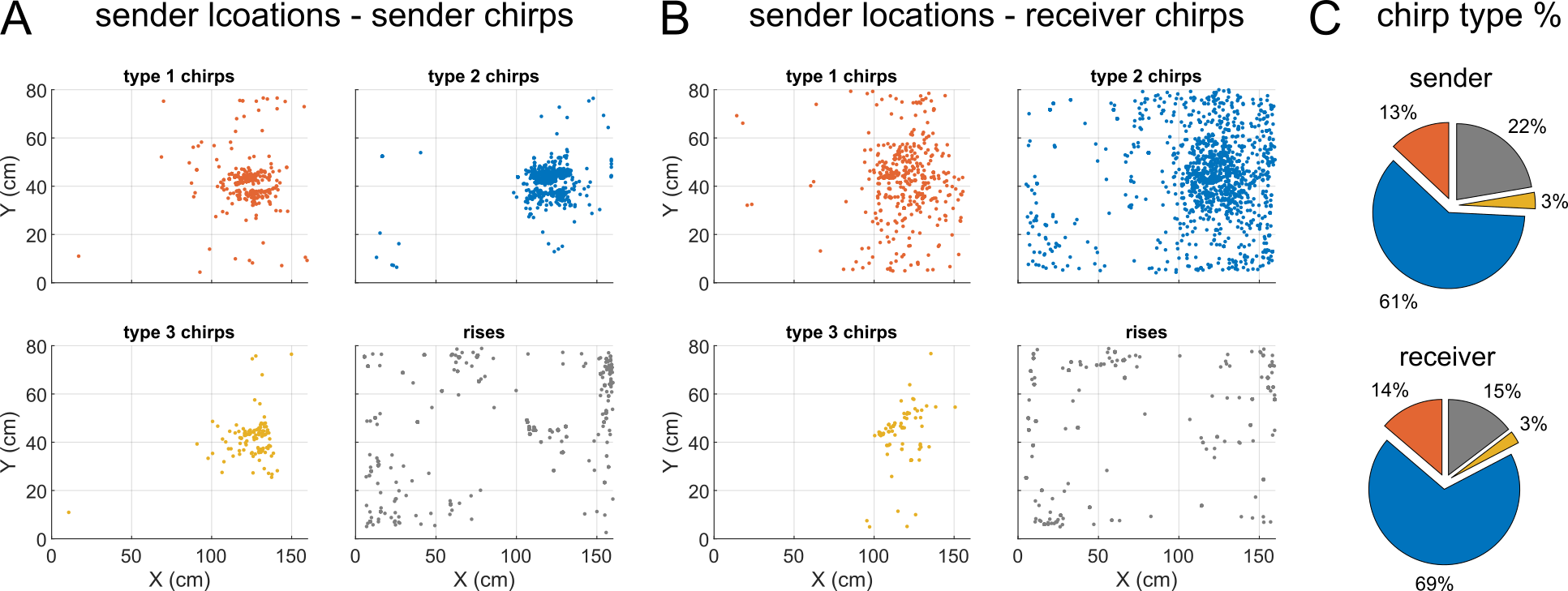

***Figure S10 Chirp type locations during novel environment explorations.*** *A: Sender fish locations during chirps of different types. Most chirps are produced while brown ghosts are swimming in close proximity to a caged conspecific (see* ***Figure 8****). This is particularly evident for freely swimming “sender” fish. Chirps produced by caged fish (“receiver”) are more widely distributed. Rises are produced when fish are perpendicularly oriented, along the wall (right) or half hidden behind shelters or plastic barriers. B: Sender fish locations during chirps produced by the caged conspecific (receiver). Receiver chirps are produced at similar locations, and in similar percentage (C).*

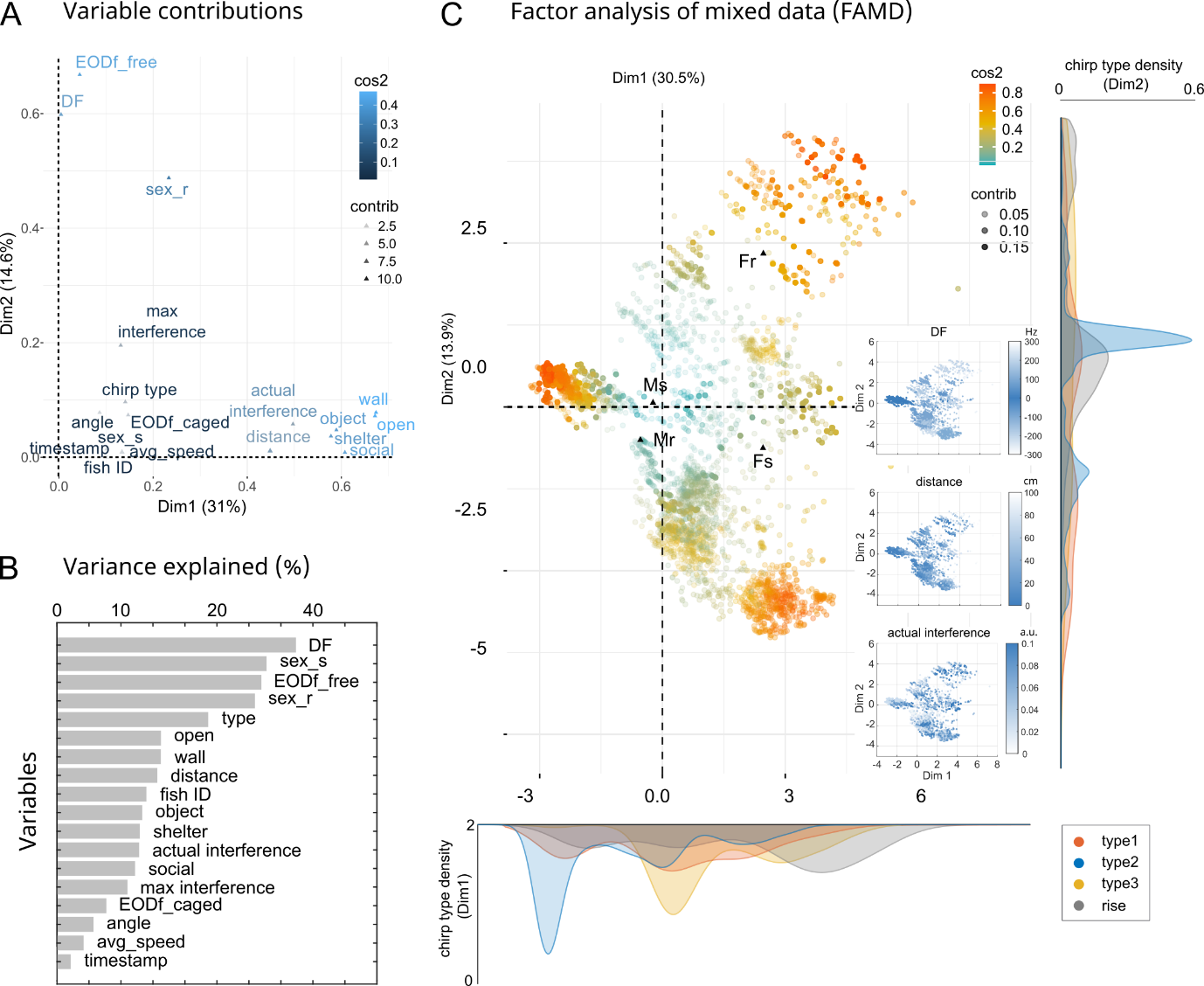

***Figure S11. Factor analysis of mixed data (FAMD) – novel environment exploration.*** *A: The scatter plot represents the contribution of chirp-related variables to the 2 main components of the transformed space. The contribution of each variable is indicated by both the position and the color of the corresponding marker on the plot. The variables closer to the origin contribute less to the overall sample variance. Variable contribution (contrib) is coded by color intensity and the quality of the representation by color hue (cos2). B: Bar plot showing the total variance (i.e. sum of each variable loading on all the 3 dimensions) explained by each variable in the transformed space. C: Representation of all 7894 chirps in the transformed coordinates. Triangles indicate the coordinates of the qualitative variable centroids (Fs = female sender, Fr = female receiver, Ms = male sender, Mr = male receiver). The contribution of individual chirps is color coded as in B. The clustering is based on both quantitative and qualitative coordinates (fish ID, sex sender, sex receiver, chirp type, average swimming speed, distance and angle between fishes, time spent in the tank ROIs, EOD frequencies of the interacting fish and DF, estimated beat interference and maximum interference possible). The marginal histograms show the kernel density distributions of different chirp types. In the insets, chirps are color-coded according to DF, distance and interference.*

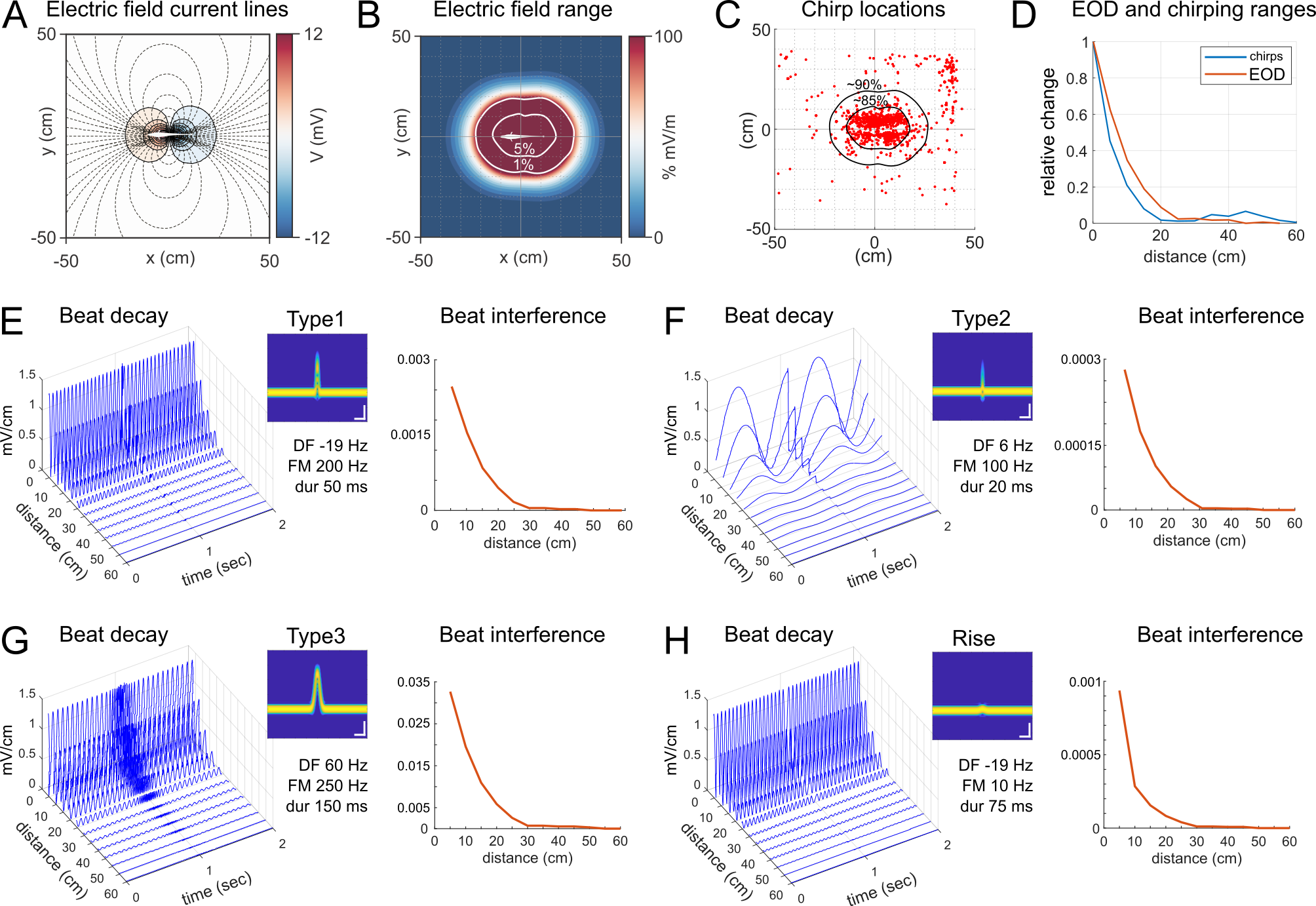

***Figure S12. Chirps interfere with the beat within the electric field range.*** *A: Electric field generated by a 16 cm 3D dipole modeling an electric fish of the same size using BEM (boundary element methods). Iso-potential lines are shown for the near-field range in different colors, based on field polarity. Current is represented by the dashed lines, perpendicular to them. B: Electric field intensity mapped around the same modeled fish. The level lines represent the 1% and 5% intensity of the electric field generated by the ideal fish. C: Scatter plot of chirp locations. The overlay is centered at the origin and corresponds to 90% and 85% of all chirps produced, respectively. D: Plot showing the intensity range of an EOD mimic calculated at 22-25 °C and 200 μS (red) and the distribution range of chirps emitted by real fish (blue), for comparison. E-H: 3D plots showing the ideal beats calculated for different sinewave pairs during different chirp types and plotted over distance. The 2D plots on the side represent the beat interference (calculated using a threshold of 1% of maximum beat amplitude) caused by each chirp type over distance. Scale bars in the spectrograms are 100 ms and 100 Hz.*
